# Redundancy principle for optimal random search in biology

**DOI:** 10.1101/210443

**Authors:** Z. Schuss, K. Basnayake, D. Holcman

## Abstract

Chemical activation rate is traditionally determined by the diffusion flux into an absorbing ball, as computed by Smoluchowski in 1916. Thus the rate is set by the mean first passage time (MFPT) of a Brownian particle to a small target. This paradigm is shifted in this manuscript to set the time scale of activation in cellular biology to the mean time of the first among many arrivals of particles at the activation site. This rate is very different from the MFPT and depends on different geometrical parameters. The shift calls for the reconsideration of physical modeling such as deterministic and stochastic chemical reactions based on the traditional forward rate, especially for fast activation processes occurring in living cells. Consequently, the biological activation time is not necessarily exponential. The new paradigm clarifies the role of population redundancy in accelerating search processes and in defining cellular-activation time scales. This is the case, for example, in cellular transduction or in the nonlinear dependence of fertilization rate on the number of spermatozoa. We conclude that statistics of the extreme set the new laws of biology, which can be very different from the physical laws derived for individuals.

## 1 Introduction

Why are specialized sensory cells so sensitive and what determines their efficiency? For example, rod photoreceptors can detect a single photon in few tens of milliseconds [1], olfactory cells sense few odorant molecules on a similar time scale, calcium ions can induce calcium release in few milliseconds in neuronal synapses, which is a key process in triggering synaptic plasticity that underlies learning and memory. Decades of research have revealed the molecular processes underlying these cellular responses, that identify molecules and their rates, but in most cases, the underlying physical scenario remains unclear.

Here, we discuss how the many molecules, which are obviously redundant in the traditional activation theory, define the *in vivo* time scale of chemical reactions. This redundancy is particulary relevant when the site of activation is physically separated from the initial position of the molecular messengers. The redundancy is often generated for the purpose of resolving the time constraint of fast-activating molecular pathways. Activation occurs by the first particle to find a small target and the time scale for this activation uses very different geometrical features (optimal paths) than the one described by the traditional mass-action law, reaction-diffusion equations, or Markov-chain representation of stochastic chemical reactions.

We also discuss the role of the fastest particle to arrive at a small target in the context of fertilization, where the enormous and disproportionate number of spermatozoa, relative to the single ovule, remains an enigma. Yet, when the number of sperms is reduced by four, infertility ensues [2]. The analysis of extreme statistics can explain the waste of resources in so many natural systems. So yes, numbers matter and wasting resources serve an optimal purpose: selecting the most fitted or the fastest.

More specifically, mass-action theory of chemical reactions between two reactants in solution, *A* and *B*, is expressed as

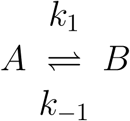

where *k*_1_ and *k*_*−1*_ are the forward and backward reaction rates, respectively. The computation of the backward rate has long history that begins with Arrhenius’ law *k*_−1_ = *Ae*^*E/kT*^ where *A* is a constant, and *E* is activation energy, then Kramers’ rate, derived from the molecular stochastic Langevin equation, which gives the prefactor *A*. For the past sixty years, chemical physicists computed the activation energy *E* and clarified the role of the energy landscape, with extensions to applications in chemistry, signal processing (time to loss of lock in phase trackers [3]), finance (time for binary option price to reach a threshold), and many more [4].

In contrast, the forward rate *k*_1_ represents the flux of three-dimensional Brownian particles arriving at a small ball or radius *a*. Smoluchowski’s 1916 forward rate computation reveals that

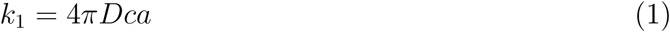

where *D* is the diffusion coefficient, when the concentration *c* is maintained constant far away from the reaction site. When the window is in a smooth surface or inside a hidden cusp, the forward rate *k*_1_ is the reciprocal of the MFPT of a Brownian particle to the window. The precise geometry of the activating small windows has been captured by general asymptotics of the mean first arrival time at high activation energy. This mean time is, indeed, sufficient to characterize the rate, because the binding process is Poissonian and the rate is precisely the reciprocal of the MFPT. These computations are summarized in the narrow escape theory [5–9]. For example, when the small absorbing window represents binding at a surface, the forward rate is given by

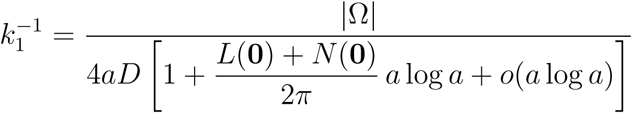

with |Ω| the volume of the domain of Brownian motion, *a* is the radius of the absorbing window [5], and *L*(0) and *N*(0) are the principal mean curvatures of the surface at the small absorbing window.

The forward rate *k*_1_ has been used in almost all representations of chemical reactions: it is the basis of the Gillespie algorithm for generating statistics of stochastic simulations with rate *k*_1_. It has also been used in coarse-graining stochastic chemical-reactions into a Markov chain. However, the rate *k*_1_ is not used in simulations of Brownian trajectories. In the latter case, the statistics of arrival times can be computed directly. Obviously, diffusion theory computes *k*_1_ from the mean arrival rate of a single particle. However, in cell biology, biochemical processes are often activated by the first (fastest) particle that reaches a small binding target, so that the average time for a single particle does not necessarily represent the time scale of activation. Thus even the Gillespie algorithm would give a rate, sampled with mean *k*_1_. But, as seen below, its statistics is different from the rate of the fastest arrival. This difference is the key to the determination of the time scale of cellular activation that can be computed from full Brownian simulations and/or asymptotics of the fastest particle. This is an important shift from the traditional paradigm.

## 2 Biochemical reactions in cell biology

Chemical activation in cell biology starts with the binding of few molecules (Fig 1). The signal is often amplified so that a molecular event is transformed into a cellular signal. How fast is this activation? What defines its time scale? when there is a separation between the site of the first activation and that of amplification. When particle move by diffusion, is it sufficient that the first particle arrives to a receptor to open it and lead to an avalanche either through the entry of ions or the opening of the neighboring receptors. Thus the time scale of activation is not given by the reciprocal of the forward rate, but rather by the extreme statistics, that is, by the mean arrival time of the first particle to the activation site (the target).

**Figure 1:**
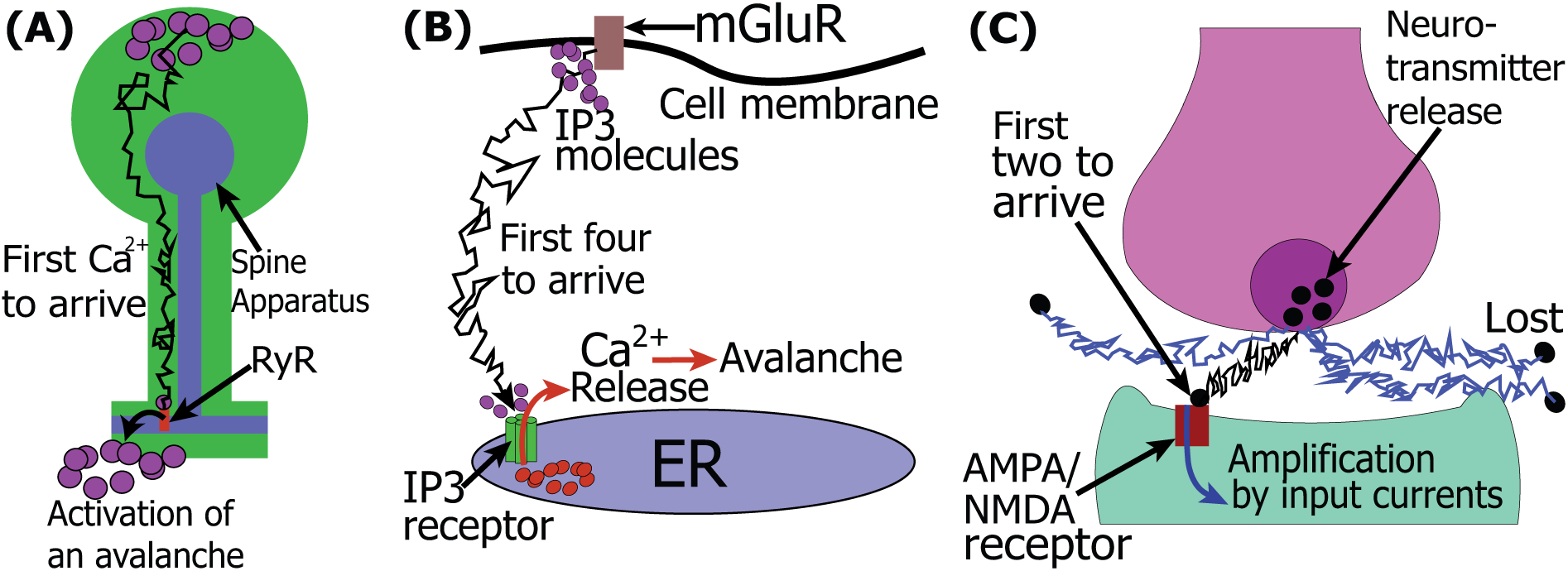
**(A)** Calcium-induced-calcium-release in a dendritic spine. The first Ryanodine Receptor (RyR) at the base of the spine apparatus that opens is triggered by the fastest calcium ion. An avalanche of calcium release ensues by opening the neighboring receptors. This leads to rapid amplification at a much shorter time than the MFPT of the diffusing calcium ions. (B) Activation of calcium release by IP3 receptors, which are calcium channels gated by IP3 molecules, which function as secondary messengers. When the first IP3 molecules arrives at the first IP3R, its calcium release induces an avalanche due to the opening of subsequent IP3 receptors. **(C)** In the post-synaptic terminal, the influx of ions due to the opening of NMDA/AMPA receptors is the amplification process. The signal of the pre-synaptic signal transmitted by the neurotransmitter molecules diffusing in the synaptic cleft and the time scale of the amplification are determined by the fastest molecules that arrive at the receptor targets and open them.

The statistics of the first particle to arrive to a target can be computed from the statistics of a single particle when they are all independent and identically distributed [10–13]. With *N* non-interacting i.i.d. Brownian trajectories (ions) in a bounded domain Ω to a binding site, the shortest arrival time *τ*^1^ is by definition

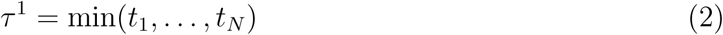

where *t*_*i*_ are the i.i.d. arrival times of the *N* ions in the medium. The narrow escape problem (NEP) is to find the PDF and the MFPT of *τ*^1^. The complementary PDF of *τ*^1^,

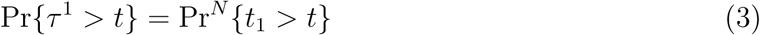

Here Pr {*t*_1_ *> t*} is the survival probability of a single particle prior to binding at the target. This probability can be computed by solving the diffusion equation

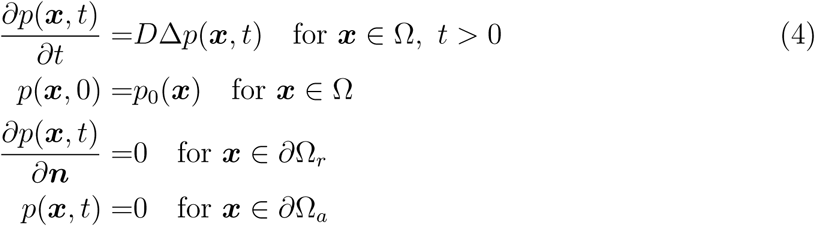

where the boundary *∂*Ω contains *N*_*R*_ binding sites 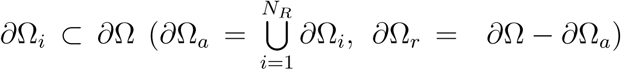. The single particle survival probability is

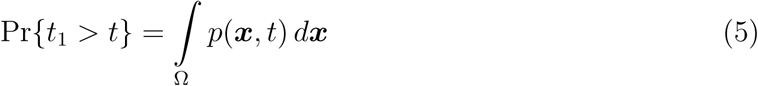

so that 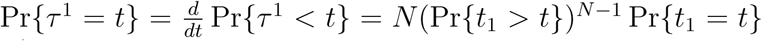, where 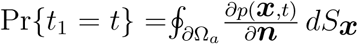 and 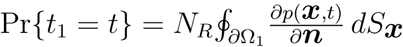. The pdf of the arrival time is

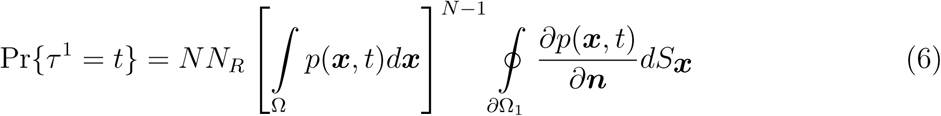

which gives the MFPT

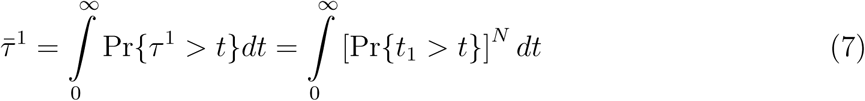

The shortest ray from the source to the absorbing window *δ*_*min*_ plays a key role, because the fastest trajectory is as close as possible to that ray. The diffusion coefficient is *D*, the number of Brownian particles is *N*, *s*_2_ = |***x – A***|, ***x*** the positions of injection, and the center of the window is ***A***. The asymptotic laws for the expected first arrival time of Brownian particles to a target for large *N*, are,

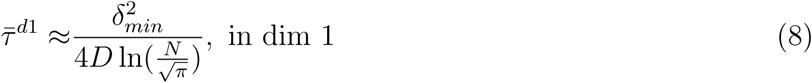

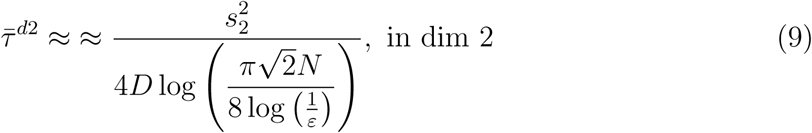

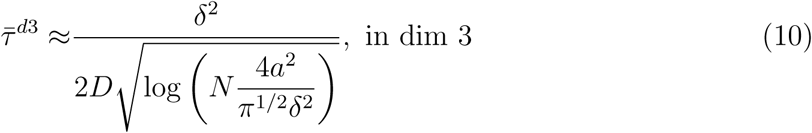

These formulas show that expected arrival time of the fastest particles is *O*(1*/* log(*N*)) (see Fig. 2). They should replace the classical forward rate in models of activation in biochemical reactions.

**Figure 2:**
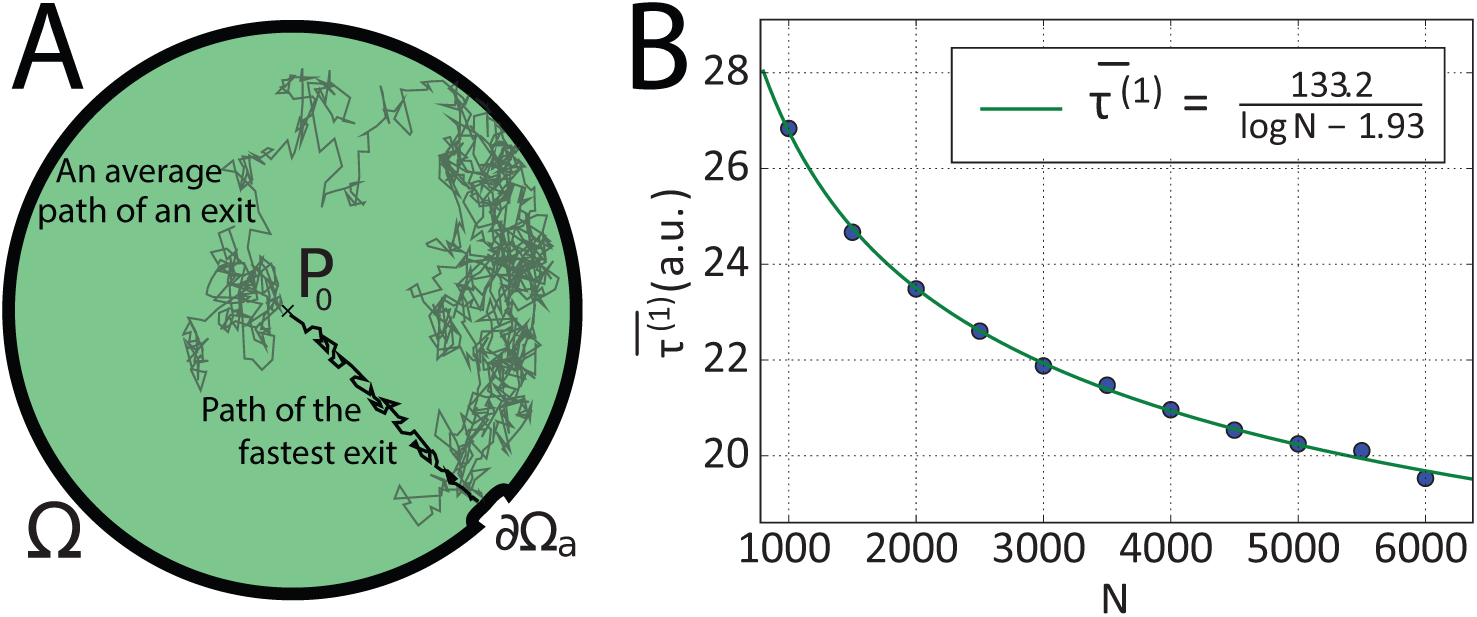
Escape through a narrow opening in a planar disk. **A.** The geometry of the NEP for the fastest particle. **B.** Plot of the expected arrival time of the fastest path vs the number of particles *N*. The asymptotic solution (red curve) is fit to 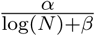.

## 3 Examples of signal transduction

There are many examples of activation or transduction by the arrival of the first particle at the activation site, which defines the time scale of [1]. First, neuronal connections often occur on a dendritic spine, where the fast calcium increase in dendrites may happen a few milliseconds after initiation in the spine head. This time scale is incompatible with diffusion alone [5]. It was shown recently that such a short time scale is generated by an avalanche reaction triggered by the arrival of the fastest calcium ion to a receptor (see Fig. 1). A similar process occurs in a fly’s photoresponse to the absorption of a single photon: the first TRP channel that opens is due to the local diffusion of several IP3 molecules and calcium ions produced after rhodopsin activation.

In another context, the post-synaptic current is generated when the first receptor of a neurotransmitter is activated (Fig. 1). This activation is mediated by the release of thousands of neurotransmitters from the pre-synaptic terminal. Two binding events on the same receptor are required for activation. The time scale of this process is defined by the arrival of the first two neurotransmitters at the same receptor. The asymmetry between the number of neurotransmitters (2000 to 3000) and the low number of receptors (5 to 50) compensates for the low probability of finding the small targets (receptors) [5].

Another transduction pathway is the one of IP3, which begins with the activation of mGluR receptors and leads to IP3 production. These molecules have to diffuse and to bind to the first IP3 receptor in order to trigger calcium release, which leads to the amplification of the response. The number of activated IP3 and the location of IP3 receptors sets the time scale of this transduction pathway.

## 4 A nonlinear effect of spermatozoa redundancy on fertility

In the key step of fertilization, sperms have to find the ovule within the short time it is fertile (Fig. 3). Thus the arrival of the first few sperms gives them much higher chance to fertilize the ovule. The trajectories of the fastest spermatozoa, which result from the directed motion model, reveal that the extreme trajectories trace the most effective path of that dynamics. These are the optimal (bang-bang) solutions of the classical control problem (Fig. 3 and [14]). This result suggests that linear trajectories might not be generated by any chemotaxis at a distance of a few centimeters, which is too far away from the source. Similarly, it is not clear how other physical scenarios, such as rheotaxis or thermotaxis contribute to sperm guidance [15]. It is thus conceivable that the extreme statistics are responsible for selection of the fastest trajectories determined by sperm dynamics in the uterus and fallopian tubes. The number of spermatozoa is thus the main determinant of the selection and since the mean time for the first one to arrive is *O*(1*/logN*), a large number of them is necessary to affect the search process. In addition, it is well documented that reducing their number by a factor of 4 may cause infertility [2].

**Figure 3:**
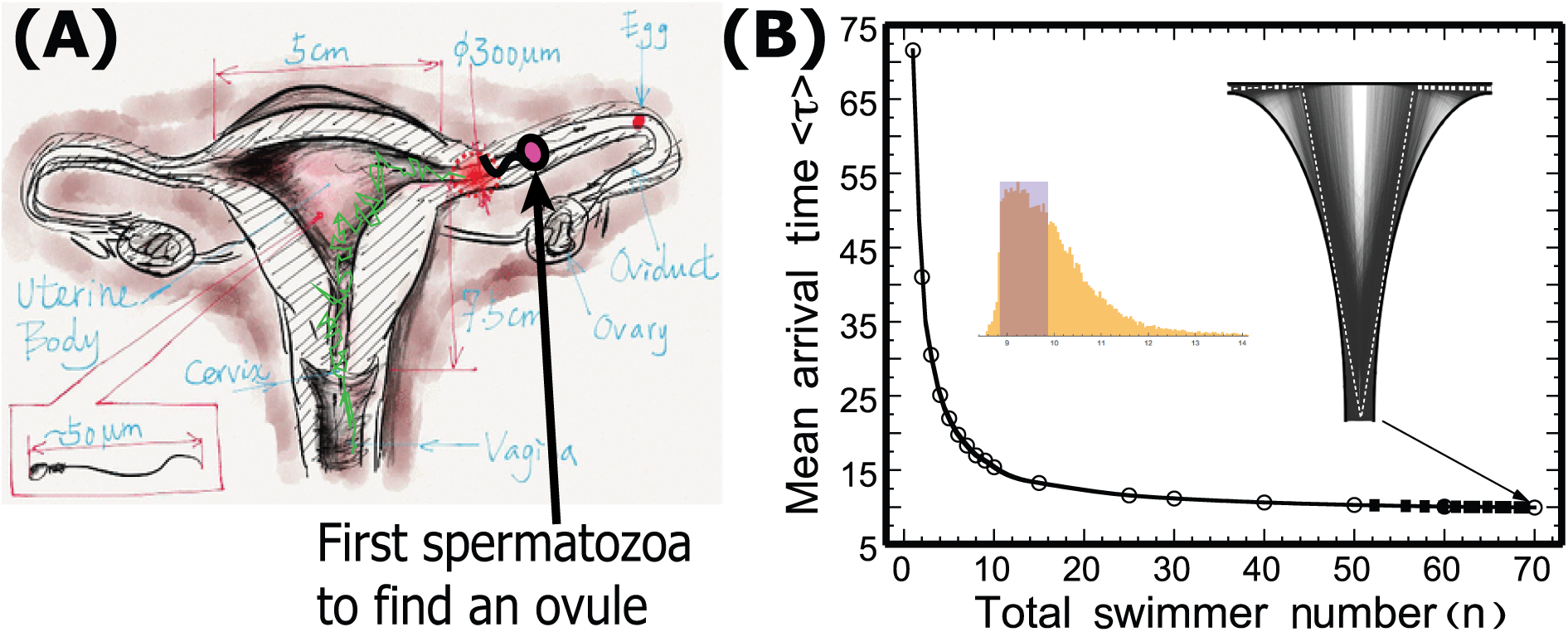
**(A)** Schematic representation of the uterus and the diffusive trajectory of the sperm that reaches an ovule. The possibility to form a zygote is determined by whether the time of the fastest sperm to reach the ovule is within the fertile period. **(B)** In a Brownian simulation in a schematic uterus, the trajectories with short arrival time are concentrated along two symmetric optimal paths (white dashed line) [14].

## 5 Morale of the story

Molecular-level simulations of activation processes should avoid the Gillespie algorithm and reaction-diffusion equations especially in the context of cellular fast activation induced by molecular pathways. They should use straightforward Brownian paths and extreme statistics, as given in the formulas above. The large number of paths produces optimal trajectories that set the observed time scale. Disproportionate numbers of particles in natural processes should not be considered wasteful, but rather, they serve a clear purpose: they are necessary for generating the fastest response. This property is universal ranging from the molecular scale to the population level. It seems that nature’s strategy for optimizing the response time is not necessarily defined by the physics of the motion of an individual particle, but rather by the collective extreme statistics.

